# clusterExperiment and RSEC: A Bioconductor package and framework for clustering of single-cell and other large gene expression datasets

**DOI:** 10.1101/280545

**Authors:** Davide Risso, Liam Purvis, Russell Fletcher, Diya Das, John Ngai, Sandrine Dudoit, Elizabeth Purdom

**Affiliations:** Division of Biostatistics and Epidemiology, Weill Cornell Medicine, New York, NY, USA; Department of Statistics, UC Berkeley, Berkeley, CA, USA; Department of Molecular and Cell Biology, UC Berkeley, Berkeley, CA, USA; Berkeley Institute for Data Science, UC Berkeley, CA USA; Division of Biostatistics, UC Berkeley, Berkeley, CA, USA

## Abstract

Clustering of genes and/or samples is a common task in gene expression analysis. The goals in clustering can vary, but an important scenario is that of finding biologically meaningful subtypes within the samples. This is an application that is particularly appropriate when there are large numbers of samples, as in many human disease studies. With the increasing popularity of single-cell transcriptome sequencing (RNA-Seq), many more controlled experiments on model organisms are similarly creating large gene expression datasets with the goal of detecting previously unknown heterogeneity within cells.

It is common in the detection of novel subtypes to run many clustering algorithms, as well as rely on subsampling and ensemble methods to improve robustness. We introduce a Bioconductor R package, `clusterExperiment`, that implements a general and flexible strategy we entitle Resampling-based Sequential Ensemble Clustering (RSEC). RSEC enables the user to easily create multiple, competing clusterings of the data based on different techniques and associated tuning parameters, including easy integration of resampling and sequential clustering, and then provides methods for consolidating the multiple clusterings into a final consensus clustering. The package is modular and allows the user to separately apply the individual components of the RSEC procedure, i.e., apply multiple clustering algorithms, create a consensus clustering or choose tuning parameters, and merge clusters. Additionally, `clusterExperiment`provides a variety of visualization tools for the clustering process, as well as methods for the identification of possible cluster signatures or biomarkers.

The package `clusterExperiment`is publicly available through the Bioconductor Project, with a detailed manual (vignette) as well as well documented help pages for each function.

## 1 Introduction

The clustering of samples or genes is one of the most common tasks in gene expression studies and, in many studies with large numbers of samples, the precise allocation of samples to clusters is critical to identify biological subtypes. With single-cell transcriptome sequencing (scRNA-Seq) studies, in particular, the boundaries between subtypes can be quite fuzzy, with individual cells on the boundary of clusters. Similarly, outlying contaminant cells are also common. These problems highlight the need for robust identification of clusters, and several clustering algorithms have been recently proposed for the specific task of clustering single-cell sequencing data [1–7].

We introduce here the Bioconductor package `clusterExperiment`, which implements not a specific clustering algorithm, but a general and flexible framework in particular useful for the clustering of cells based on single-cell RNA-Seq data. We also introduce a specific clustering workflow entitled Resampling-based Sequential Ensemble Clustering (RSEC). It enables researchers to easily try a variety of different clustering algorithms and associated tuning parameters and generate a stable consensus clustering from these many candidate clusterings. Specifically, given user-supplied base clustering algorithms and associated tuning parameters, RSEC runs the algorithms and generates the corresponding collection of candidate clusterings, with the option of resampling cells and of using a sequential clustering procedure as in [8]. Resampling greatly improves the stability of clusters and has been frequently suggested in gene expression clustering [9] and for single-cell studies specifically [1]. Additionally, considering an ensemble of methods and tuning parameters allows to capitalize on the different strengths of the base algorithms and avoid the subjective selection of tuning parameters. RSEC provides a strategy for defining a consensus clustering from the many candidate clusterings and a method of further merging similar clusters that do not show strong individual gene expression differences.

Unlike many existing clustering software for single-cell sequencing and gene expression data, `clusterExperiment` provides a flexible framework that allows for user customization of the clustering algorithm and accompanying manipulation of the data. Finally, the `clusterExperiment`package is fully integrated into the Bioconductor software suite, inheriting from the existing `SingleCellExperiment` class (a baseline class for storing single-cell data) [10], and interfaces with common differential expression (DE) packages like `limma` [11], `MAST` [12], and `edgeR` [13] to find marker genes for the clusters.

## 2 Design and Implementation

In what follows, we refer to a “clustering” as the set of clusters found by a single run of a clustering method, while a “cluster” refers to a set of samples within a clustering.

### 2.1 RSEC Workflow

The `clusterExperiment`package provides a novel workflow for creating a unified clustering from many clustering results which we entitle Resampling-based Sequential Ensemble Clustering (RSEC). RSEC formalizes many choices that are often seen in practice when clustering large RNA-Seq expression datasets. In particular, RSEC formalizes the process of manually experimenting with many different parameter choices by systematically running them all and then provides a formal mechanism for creating an ensemble or consensus clustering from the results.

The RSEC workflow comprises the following steps, demonstrated in Fig 1:

**Fig 1.**
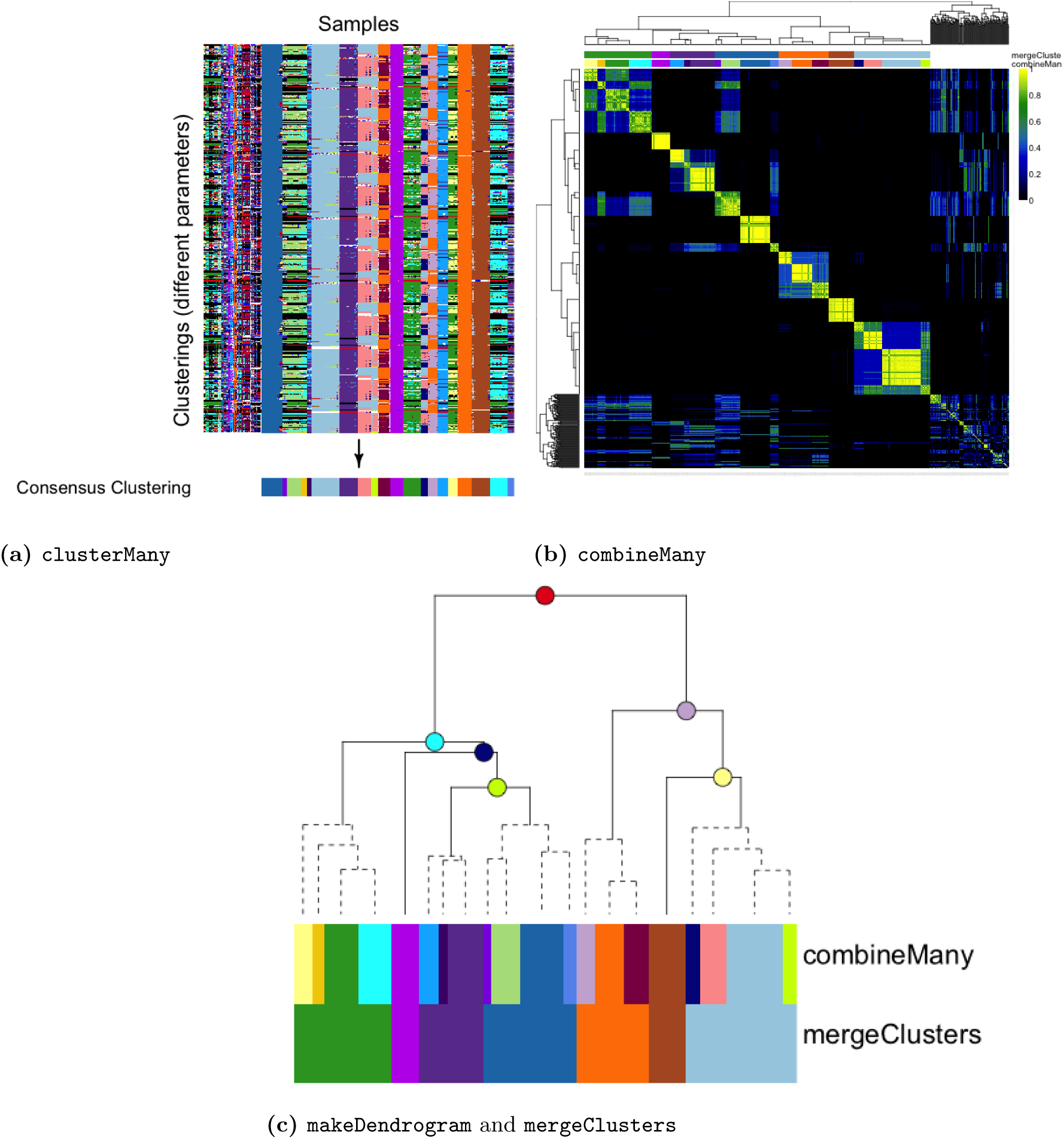
*Main steps of RSEC workflow, demonstrated on the olfactory epithelium dataset.* (a) The clusterMany step produces many clusterings from the different combinations of algorithms and tuning parameters. These clusterings are displayed using the plotClusters function. Each column of the plot corresponds to a sample and each row to a clustering from the clusterMany step. The samples in each row are color-coded by their cluster assignment in that clustering; samples that are not assigned to a cluster are left white. The colors across different clusterings (rows) are assigned so as to have similar colors for clusters with similar samples across clusterings. The consensus clustering obtained from the combineMany step is also shown below the individual clusterings. (b) The combineMany step finds a consensus clustering across the clusterMany clusterings based on the co-occurrence of samples in these clusterings. The heatmap of the matrix of co-occurrence proportions is plotted using the plotCoClustering function. The resulting cluster assignments from combineMany are color-coded above the matrix, as are the assignments from the next step, mergeClusters. (c) The makeDendrogram step creates a hierarchy between the consensus clusters and then similar clusters in sister nodes are merged with mergeClusters. Plotted here with the function plotDendrogram is the hierarchy of the clusters from makeDendrogram, with merged nodes indicated with dashed lines. The combineMany clusters and resulting mergeClusters clusters are indicated as color-coded blocks on the right of the dendrogram, sized according to the number of samples in the clusters. The estimated proportions of DE genes of each node are shown in Supplementary Fig S1b.

1. `clusterMany` Implementation of one or more clustering methods across a wide range of tuning parameter and data dimensionality choices.
2. `combineMany` Determination of a single consensus clustering from these many clusterings.
3. Merging of individual clusters from the consensus clustering, involving two steps:
  - `makeDendrogram` Defining a hierarchical clustering of the clusters,
  - `mergeClusters` Merging clusters along that hierarchy by collapsing into a single cluster sister nodes between which only a small proportion of genes show differential expression.

The main `RSEC` wrapper function in the `clusterExperiment`package is our recommended implementation of this workflow. In addition to performing all workflow steps within a single function, it implements our preferred choice of subsampling and sequential clustering in the `clusterMany` step. It is our experience that these choices are particularly relevant for single-cell sequencing and other large RNA-Seq experiments. However, the user has the option to perform the individual steps separately using a sequence of function calls and make more customized choices within this workflow. The intermediate clusterings found at all of these steps are retained using a dedicated class for RSEC clustering results, so that the user can examine the impact of each step using the visualization tools of the `clusterExperiment`package.

### 2.2 Clustering Procedures

We briefly describe here the core procedures that make up the RSEC workflow. More details can be found in the Supplementary Text.

#### 2.2.1 Generating Many Clusterings (clusterMany)

The function `clusterMany` allows the user to easily select a range of clustering algorithms and tuning parameters and create different clusterings for each combination thereof. The option is given to parallelize computations across multiple cores. A few examples of parameters that can be compared are: the dimensionality reduction method, the number of dimensions, the number of clusters *K* (if appropriate for the clustering method), whether to subsample to create the clustering, and whether to sequentially detect clusters.

The choices regarding whether to subsample the data and whether to sequentially detect clusters can be paired with any clustering algorithm, but their implementation can be non-trivial, which makes the implementation of these procedures, independent of a particular clustering method, particularly useful.

##### Subsampling

The subsampling option in `clusterExperiment`generates clusterings based on randomly sampled subsets of the full set of *n* observations, and each resampled dataset is clustered with the baseline clustering algorithm. Subsampling defines a dissimilarity matrix between samples, with entries *D*_*ij*_ = 1 *− p*_*ij*_, where *p*_*ij*_ is defined as the proportion of times the pair of samples *i* and *j* were in the same cluster across all of the resampled datasets. Subsampling does not itself define a clustering of the samples; RSEC uses the dissimilarity matrix *D* to cluster the samples. The clustering algorithm applied to *D* does not need to be that which was used on the resampled datasets. Furthermore, because *D* is a dissimilarity matrix, with entries on a well defined scale of 0–1, it is intuitive to cluster *D* so as to constrain the level of between-sample dissimilarity within clusters [8], rather than set a particular *K* for the number of clusters, a feature which we also allow.

##### Sequential Clustering

Sequential clustering refers to the iterative detection and removal of single clusters and relies on a base clustering algorithm, like subsampling, which is iteratively re-applied after each removal of a cluster. For each clustering iteration, the sequential algorithm requires a method for specifying which is the “best” cluster so that it can be removed and the iteration continued. Our implementation of the sequential detection of clusters follows that of the tight clustering algorithm [8], but we have generalized it to fit arbitrary clustering techniques. Specifically, the “best” cluster is chosen by [8] to be that cluster which varies the least in its membership as the parameter *K* for the number of clusters is increased, as measured by the maximal percentage overlap of clusters from clusterings from *K* and *K* + 1 (the ratio of cardinality of intersection to cardinality of union).

The sequential algorithm can be particularly helpful when there is an outlying cluster that is widely different from others. Removing this cluster and re-clustering can minimize its effect on the global clustering results. Similarly, not clustering all samples can be beneficial in finding homogeneous clusters if the data are noisy or there are many samples on the boundary between clusters. (See also the clustering algorithm `pcaReduce` proposed by [2] for single-cell data, which adopts a sequential discovery of clusters, but is more narrowly based on reducing the number of dimensions for principal component analysis.)

#### 2.2.2 Creating a Consensus Clustering (combineMany)

After running `clusterMany`, the resulting `ClusterExperiment` object contains many clusterings, and the next step is to find a single clustering that represents the commonality across the many clusterings. The function `combineMany` does this by creating a dissimilarity matrix between samples, defined by entries *D*_*ij*_ = 1 *− p*_*ij*_, where *p*_*ij*_ is the proportion of clusterings for which the pair of samples *i* and *j* are in the same cluster (Fig 1b). This dissimilarity matrix is similar to that of subsampling and is likewise clustered to create an consensus clustering.

#### 2.2.3 Merging Clusters Based on a Cluster Hierarchy (mergeClusters)

The strategy of finding an consensus clustering used by `combineMany` emphasizes shared assignments across clusterings and can result in many small clusters. The number of final clusters can be adjusted in earlier steps in the clustering, but it is more intuitive in practice to visualize clustering results (e.g., using heatmaps) to see which clusters have clear differences in the expression of individual genes.

Since these individual gene effects are of great interest to practitioners and are used to evaluate the quality of a cluster, we formalize this practice by systematically evaluating the estimated number of genes with large effects between the clusters and using this as a metric to merge together clusters. We do not compare all pairs of clusters, but instead hierarchically order the clusters from `combineMany` via hierarchical clustering on the median value of each gene in each cluster. For each node in the resulting dendrogram, their children nodes define two sets of samples that are candidates for being merged into a single cluster, and each gene is individually tested for differential expression between the samples in the two sets (Fig 1c). Using these individual gene results, `mergeClusters` calculates an estimate of the proportion of differentially expressed genes at each node comparison (with multiple published methods for doing so available to the user [14–18]). These estimates provide the basis for whether to merge clusters, working from the leaves (clusters) upward.

We note that, in addition to merging clusters, the hierarchy between clusters is convenient for visualization of the clusters, as we discuss below (Section 2.4).

### 2.3 Biomarker Detection

A common task in clustering of gene expression datasets is to identify biomarkers, i.e., genes that strongly differentiate the clusters. Differential expression techniques involving hypothesis testing are often used for these tasks [11–13, 19], though it should be emphasized that such tools must be used merely for exploratory purposes, since there is severe overfitting of the data when the groups being compared have been found by clustering the data.

Since clustering of large gene expression datasets, such as single-cell RNA-Seq datasets, generally results in a large number of clusters, finding biomarkers for the clusters corresponds to testing for differential expression between many groups. A standard *F*-statistic from an ANOVA analysis is commonly used to assess differences between the groups. However, generally vast numbers of genes will have “significant” *F*-statistics and the largest *F*-statistics can easily be dominated by genes that differentiate the single, most outlying cluster (Fig. 2a). A better approach is to test for specific differences between groups, e.g., the pairwise differences between two groups, by forming contrasts from the full ANOVA model, which most DE packages allow. This has the added advantage of using all of the data for estimation of the variance parameters regardless of the size of the clusters.

**Fig 2.**
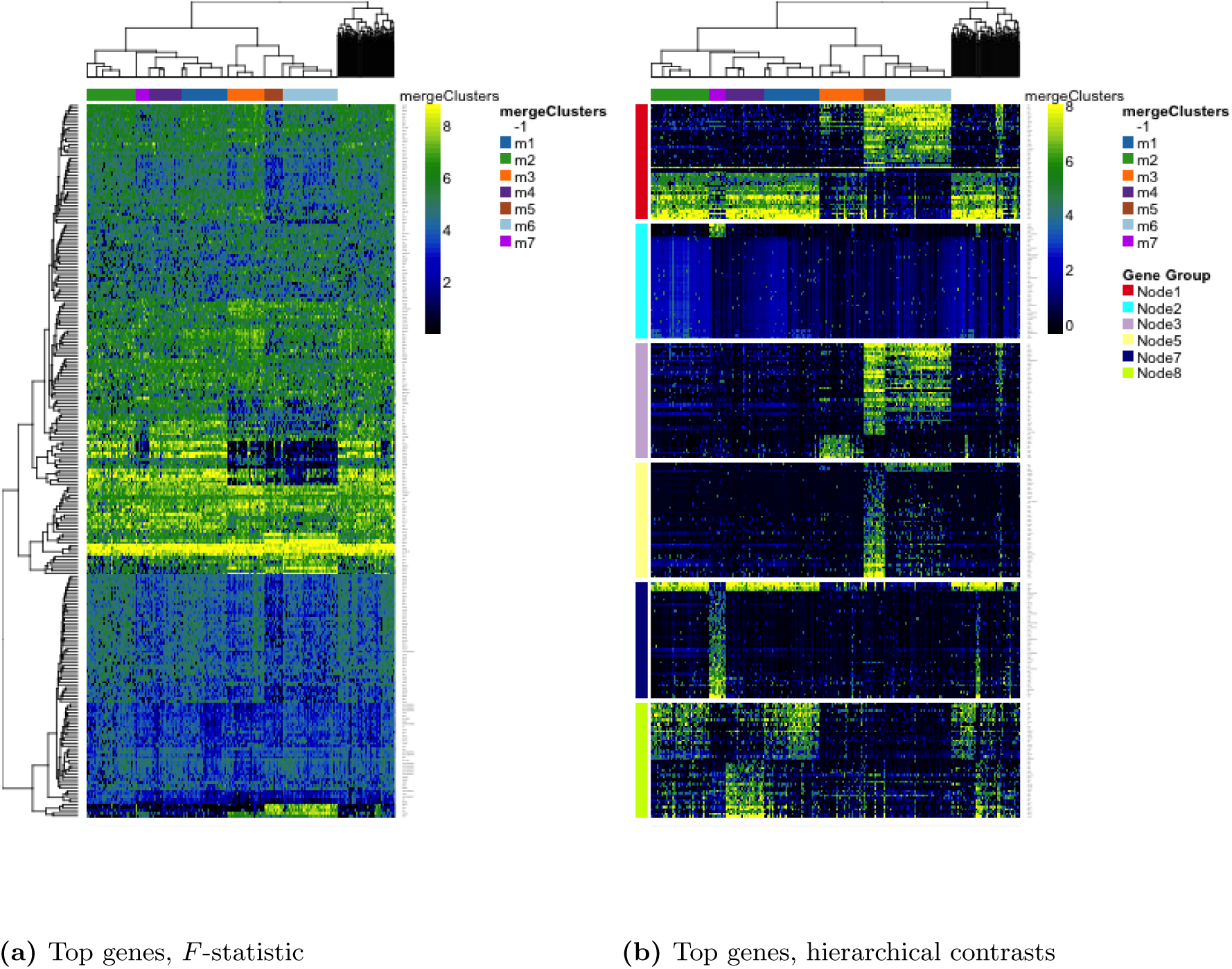
*Biomarker detection, demonstrated on the olfactory epithelium dataset.* Heatmap from the plotHeatmap function showing genes found differentially expressed (DE) between clusters by the function getBestFeatures, using both the global *F*-statistic (a) and hierarchical contrasts (b) options. Each of the contrasts in (b) corresponds to nodes in the dendrogram color-coded as in Fig 1c; we retained only the top 50 DE genes per node. Genes found DE in multiple contrasts may be plotted multiple times. For comparison purposes, in (a), we retained the top 256 DE genes according to a global *F*-statistic, where 256 is the number of unique genes shown for the hierarchical contrasts in (b).

However, a large number of clusters will imply a large set of contrasts (for example, all pairwise contrasts). Such a large number of contrasts are practically quite difficult to construct with the common DE packages. The `clusterExperimentpackage` provides tools to create relevant contrasts and optionally feeds them to the limma/voom [11, 20] package to return the relevant biomarkers. We also provide the ability to port these contrasts to `edgeR` [13] or `MAST` [12].

We provide three different kinds of contrasts that are useful for finding biomarkers for clusters:

1. all pair-wise comparisons;
2. each cluster versus all remaining clusters comparisons;
3. hierarchical comparisons, where, for each node in the cluster dendrogram from `makeDendrogram`, a comparison is performed between the two children nodes.

The last type of contrasts – hierarchical – is our novel and preferred approach for specifying comparisons based on the hierarchical relationships between the clusters, as described in the makeDendrogram step above. Unlike the other comparisons that `clusterExperiment`implements, the hierarchical comparisons allow a multi-resolution approach, identifying both biomarkers for large differences between sets of clusters and biomarkers that separate very similar clusters.

### 2.4 Visualization

The `clusterExperiment`package provides many visualization tools in addition to the above algorithmic functions, often making use of existing R and Bioconductor functionality. `clusterExperiment`brings them together in one package and, most importantly, makes it easy to integrate the clustering results with the visualization of the gene expression data.

For example, one of the most frequently used visualization methods in gene expression studies is the heatmap or pseudo-color image representation of the genes-by-samples expression matrix. `clusterExperiment`provides a function `plotHeatmap`, that relies on the `aheatmap` function in the package NMF [21] and adapts it to be specific to the results of `clusterExperiment` by optionally plotting the clustering results alongside the samples. Furthermore, a common problem in standard heatmaps is that the hierarchical clustering of the samples with a basic hierarchical clustering algorithm does not exactly correspond to the clustering found by other methods. `clusterExperiment`uses instead a hierarchical clustering of the *clusters* (described in Section 2.2.3) that keeps the clusters together and, importantly, does so by also placing the most similar clusters close to each other.

Another example is the function `plotClusters` for visually comparing large numbers of clusterings, following the work of [22]. This function calculates an alignment of the cluster assignments across samples, along with an ordering of samples, so that the user can visualize the similarity in cluster assignments (see Fig. 1a).

Other visualization tools include plotting of the hierarchy of the clusterings (`plotDendrogram`), plotting the concordance of two clusterings via a barplot (`plotBarplot`), plotting a two-dimensional representation of the data color-coded by cluster (`plotDimReduce`), and plotting boxplots of the expression levels of an individual gene per cluster (`plotFeatureBoxplot`). These are all common visualization tools for which the `clusterExperiment`package implementation makes it simple to integrate the clustering information and are demonstrated in the vignette that accompanies the package.

## 3 Results

We demonstrate the usage of the `clusterExperiment`package with a single-cell RNA-Seq dataset on neuronal stem cell differentiation in the mouse olfactory epithelium (OE) [23]. The OE is made of four major mature cell types and two progenitor cell populations. Fletcher et al. [23] used RSEC to find 13 experimentally validated clusters that clearly correspond to the known mature and progenitor cell types. The RSEC clusters were used as a starting point to identify the lineage trajectories that produce the major cell types in the OE.

Here, we independently run the entire RSEC workflow on the OE dataset with the function `RSEC`, which implements our preferred pipeline of subsampling and sequential detection of clusters across many parameter choices. We follow [23] in preprocessing the data, including filtering poor quality cells and lowly-expressed genes (see Supplementary Text, Section S-7). However, note that some of the options of RSEC differ from those used in the original article; in particular, we run far more clusterings than a typical use case in order to demonstrate the various parameters that can be varied. The following parameters were varied in the `clusterMany` step: the number of dimensions used for principal component analysis (PCA), the required similarity (*α*) from clustering of the *D* matrix from the subsampling step, the required stability of clusters in the sequential step (*β*), the starting number of clusters (*k*_0_) in the sequential search for stable clusters, and the minimum cluster size (see Table 1). This resulted in 432 separate clusterings.

**Table 1.**
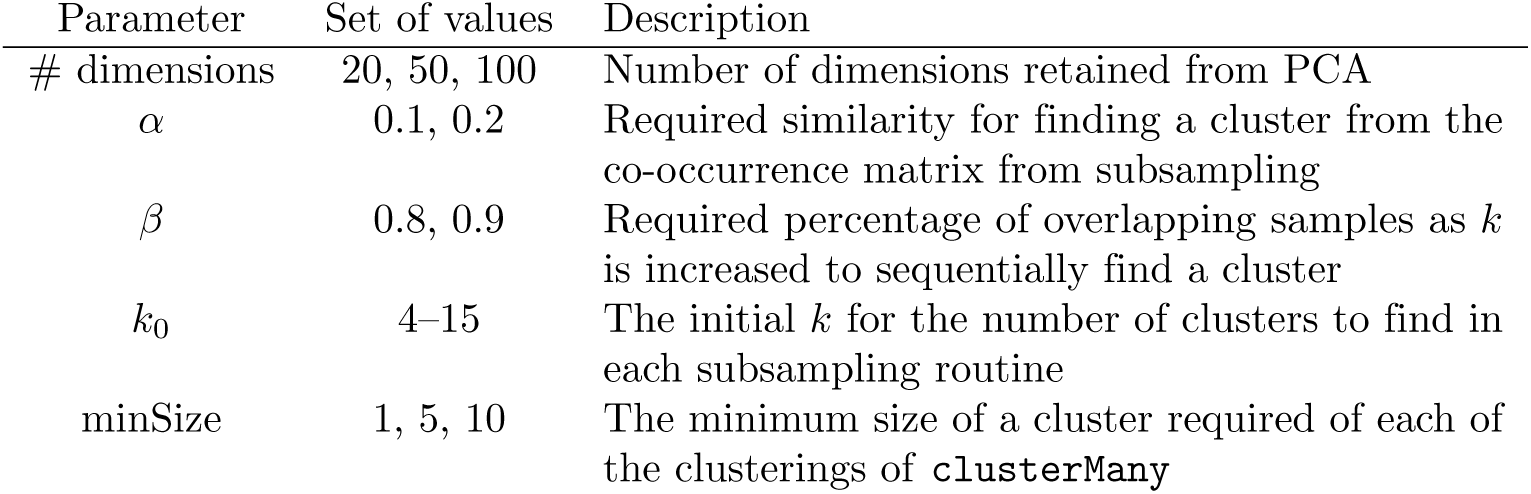
*Parameters varied when applying `RSEC` to the olfactory epithelium dataset.*

We visualize the clustering results from the `clusterMany` step using the function `plotClusters` in Fig 1a. We observe that the many different clusterings from the `clusterMany` step generally create similar clusters for the vast majority of the cells, while there exist fewer cells that are noisily assigned to different clusters as the parameters change in different clusterings. This pattern, typical for single-cell sequencing data when making use of both the subsampling and sequential steps, demonstrates the robustness provided by these approaches.

We can visualize how stable different cells were in their clusterings with a heatmap of the co-occurrence matrix (Fig. 1b): each entry of the matrix corresponds to the proportion of clusterings in which a pair of samples was clustered together. We can see strong blocks of cells that are almost always clustered together. We also note that the underlying similarity between the clusterings, despite large changes in tuning parameters, also makes the use of consensus across these clustering a logical choice.

This co-occurrence matrix forms the basis for creating a consensus between the different clusterings using the function `combineMany`, shown in Fig 1a. We can see that not all of the cells are assigned to a cluster by `combineMany`. These are samples that are either not consistently co-clustered with other samples across the different clusterings or are unassigned in a high percentage of the clusterings.

We further merge together similar clusters from the consensus clustering using `mergeClusters`, as described in Section 2.2.3. Specifically, `mergeClusters` creates a hierarchy between clusters and scores each node based on the percentage of genes showing differential expression between samples in the children nodes. Fig 1c visualizes the hierarchy found by `combineMany` as well as which clusters were merged by `mergeClusters`. We use a simple measure to estimate the proportion of genes that are differentially expressed between children node: the proportion of genes called significant at nominal level 0.05 based on the Benjamini and Hochberg [ADD REF] procedure for controlling the false discovery rate (FDR); a comparison of the different implemented methods are shown in Supplementary Fig S1b.

We would also note that `clusterExperimentmakes` it easy to switch from the merged clusters back to the consensus clusters found by `combineMany`, effectively allowing the user to explore the data at two different levels of resolution.

We finally use the function `getBestFeatures` to find genes that show strong differences in expression between clusters. For each gene, we compute DE statistics for contrasts based on the hierarchy of the tree, as well as a global *F*-statistic. The heatmaps for the top DE genes resulting from both of these approaches are shown in Fig 2 and illustrate that differences between the clusters are much more striking with the hierarchical contrasts, as compared to a global *F*-statistic.

Fig 2 also demonstrates `clusterExperiment`’s heatmap function, `plotHeatmap`, that seamlessly incorporates the clustering information into the heatmap visualization. The function automatically adds the cluster identifications of individual cells to the heatmap, but also makes use of the hierarchy that is created between the clusters (Fig. 1c). In this way, the cluster structure will be respected in the ordering of the samples, unlike a standard hierarchical clustering of the cells, yet ensures that similar clusters will be plotted close to each other rather than in an arbitrary order.

### Other Comparisons via clusterExperiment

We can also use the `clusterExperiment`package for straightforward implementation and comparison of specific clustering algorithms and parameters; the `ClusterExperiment` framework makes it easy to store and compare the results. Fig 3 shows several such examples of comparisons available in `clusterExperiment`: varying the choice of *K* for partitions around medoids (PAM), comparing 4 standard clustering functions, and comparing choices of how to calculate distances between genes. All of these different choices can be made by a choice of arguments to `clusterMany`.

**Fig 3.**
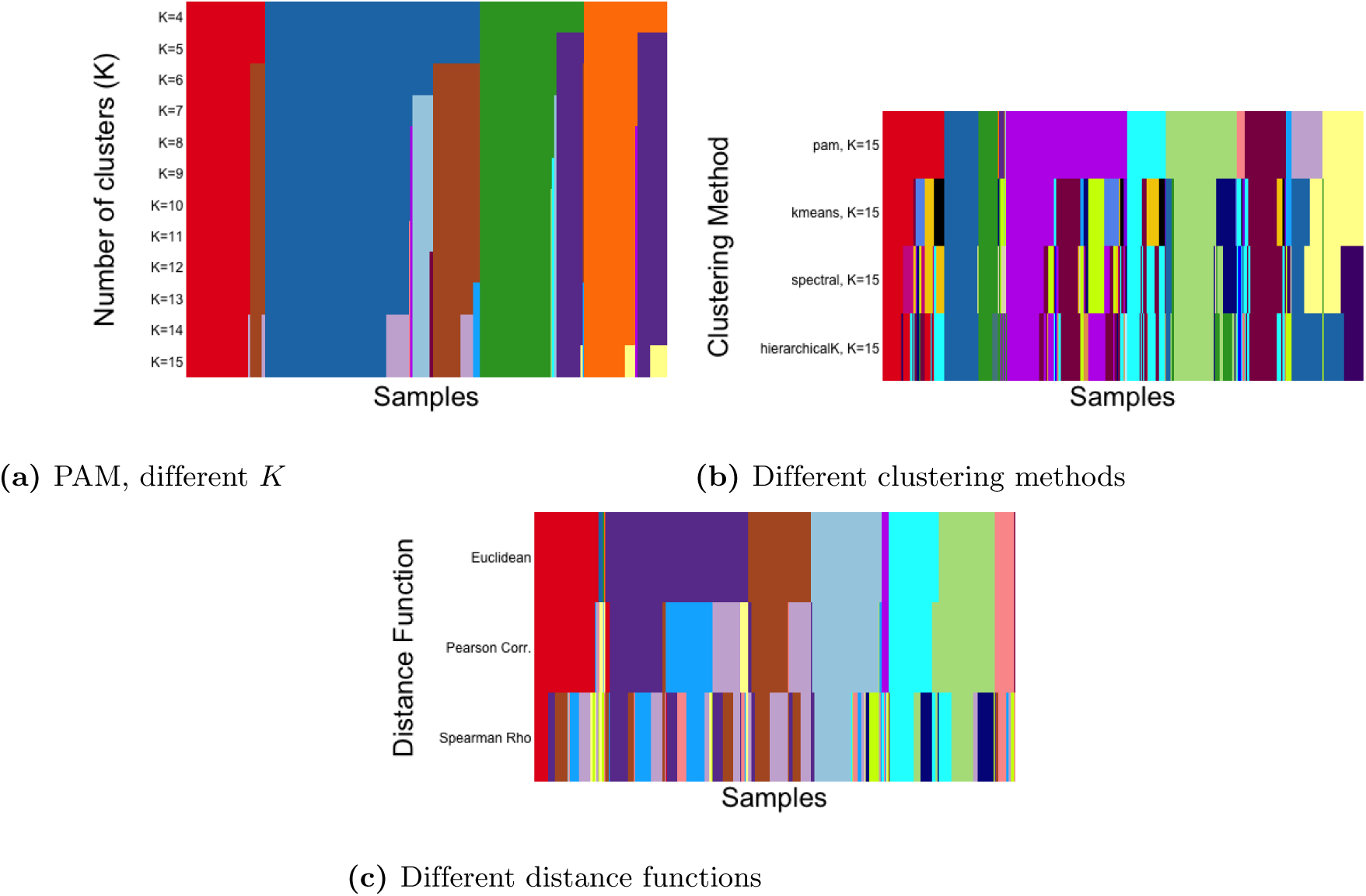
*Comparison of methods and tuning parameter choices using clusterMany and plotClusters, demonstrated on the olfactory epithelium dataset.* The figure provides examples of using clusterExperimentto compare clustering methods and tuning parameter choices via the function clusterMany to implement the clustering procedures and the function plotClusters to visualize results. (a) shows the clustering results after running PAM with different choices of *K*, the number of clusters. (b) shows the clustering results for different choices of clustering method. Each method is shown with the “best” choice of *K*, as determined by the maximum average silhouette width. (c) shows the clustering results for different between-sample distance measures. ‘Euclidean’ refers to the standard Euclidean distance; ‘Pearson Corr.’ and ‘Spearman’s Rho’ to a correlation-based distance, *d*(*i, j*) = 1/2(1*−ρ*(*i, j*)), where *ρ*(*i, j*) is either the standard Pearson correlation coefficient or the robust Spearman rank correlation coefficient between samples *i* and *j*, respectively. The clusterings in (a) and (b) were run with the top 50 PCA dimensions as input. The clusterings in (c) involve comparing different between-gene distance measures and therefore were run directly on the gene expression measures after filtering to the top 1,000 most variable genes, as determined by the median absolute deviation (MAD), a robust version of variance.

### Computational Cost

The RSEC analysis described above performed 432 clusterings on 747 samples, where each clustering consisted of an iterative sequential analysis, each iteration of the sequential clustering involved 100 clusterings of subsamples of the data, and each base clustering used 10 random restarts of the *k*-means algorithm. This analysis required a total of 4.1 hours to compute when parallelized across 32 cores on a AMD Opteron(TM) Processor 6272 node with 270GB of RAM, for a total of 106.0 hours of CPU time across all processes. The maximum amount of total combined memory across the 32 cores was 142G. This computation used the default implementation in the `clusterExperiment`package, which stores in memory an *n* × *n* matrix from each of the 100 subsamples, where *n* is the number of samples, which can result in large memory usage for the larger datasets which are now common in single-cell sequencing. The `clusterExperiment`package also provides an option to run the analysis with a more efficient memory usage for the subsampling, which results in corresponding increase in compute time (see Supplementary Text, Section S-2.1). For the OE dataset, choosing this option reduced the maximal memory usage by almost half (to 79G) and required 2,014 CPU hours to compute (again these numbers are reported as totals across the parallelization to 32 cores).

We would note that this a much larger number of clusterings that would often be performed in practice. Particularly, for very large single-cell sequencing studies, fewer than 50 clusterings would normally be sufficient given the computational costs.

## 4 Conclusion

We have demonstrated the use of a new software framework for clustering and visualizing large gene expression datasets, in particular, single-cell RNA-Seq datasets. The `clusterExperiment`package implements a wide range of clustering routines appropriate for this type of data, as well as encoding a novel workflow strategy that emphasizes robust clustering. The entire workflow is flexible and extendable by the user to other clustering routines. Furthermore, users can create clusterings externally from the package functions and upload the clustering results to a `ClusterExperiment` object to make use of the visualization and comparisons capabilities of the package.

The package `clusterExperiment`is publicly available through the Bioconductor Project, with a detailed manual (vignette) as well as well documented help pages for each function. The code for implementing all of the analyses shown here is available on the GitHub repository: www.github.com/epurdom/RSECPaper. The analysis in this paper was run using the `clusterExperiment`package version 1.99.0, R version 3.4.3, and Bioconductor 3.6.

## Supporting Information

**S1 Supplementary.pdf S1_Text.pdf** Supplemental text giving a more detailed description of the RSEC framework and its implementation, as well as information about the processing of the OE dataset.

